## Supplemental Figures and Tables for "Autism candidate gene *rbm-26* (*RBM26/27*) regulates MALS-1 to protect against mitochondrial dysfunction and axon degeneration during neurodevelopment"

### Slide 1
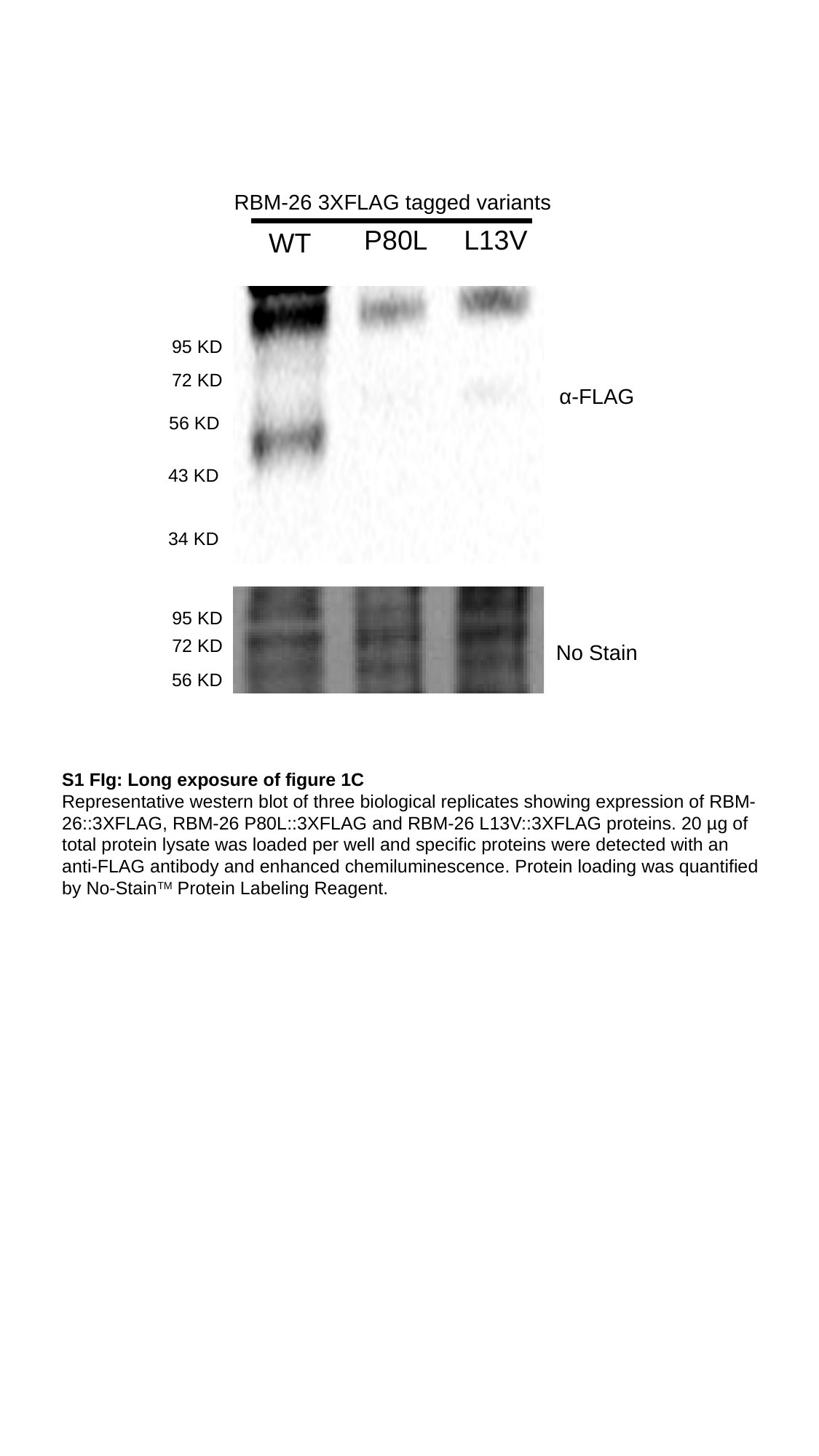

RBM-26 3XFLAG tagged variants
P80L
L13V
WT
95 KD
72 KD
α-FLAG
56 KD
43 KD
34 KD
95 KD
72 KD
No Stain
56 KD
S1 FIg: Long exposure of figure 1C
Representative western blot of three biological replicates showing expression of RBM-26::3XFLAG, RBM-26 P80L::3XFLAG and RBM-26 L13V::3XFLAG proteins. 20 µg of total protein lysate was loaded per well and specific proteins were detected with an anti-FLAG antibody and enhanced chemiluminescence. Protein loading was quantified by No-Stain^TM^ Protein Labeling Reagent.

### Slide 2
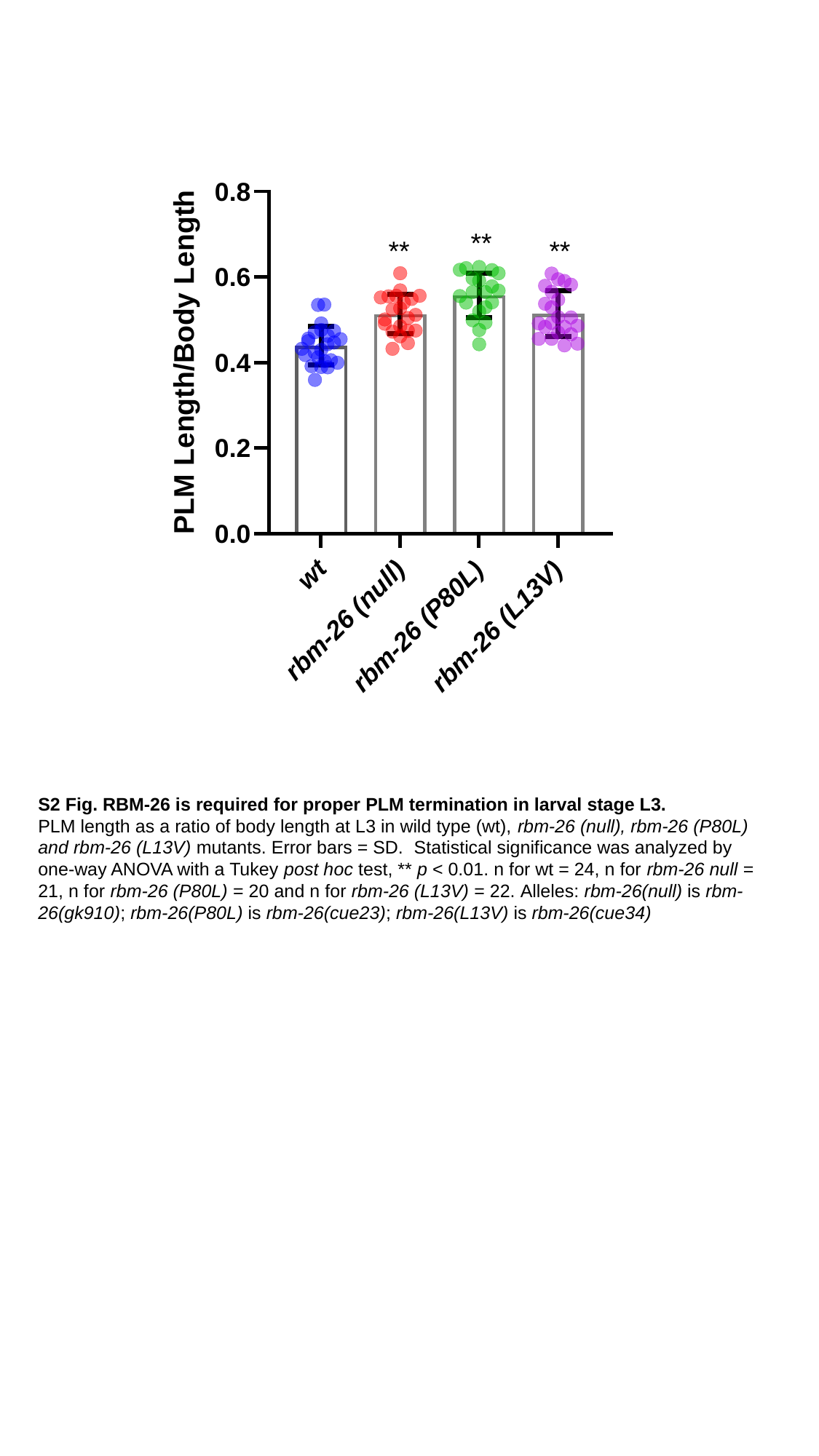

**
**
**
S2 Fig. RBM-26 is required for proper PLM termination in larval stage L3.
PLM length as a ratio of body length at L3 in wild type (wt), rbm-26 (null), rbm-26 (P80L) and rbm-26 (L13V) mutants. Error bars = SD.  Statistical significance was analyzed by one-way ANOVA with a Tukey post hoc test, ** p < 0.01. n for wt = 24, n for rbm-26 null = 21, n for rbm-26 (P80L) = 20 and n for rbm-26 (L13V) = 22. Alleles: rbm-26(null) is rbm-26(gk910); rbm-26(P80L) is rbm-26(cue23); rbm-26(L13V) is rbm-26(cue34)

### Slide 3
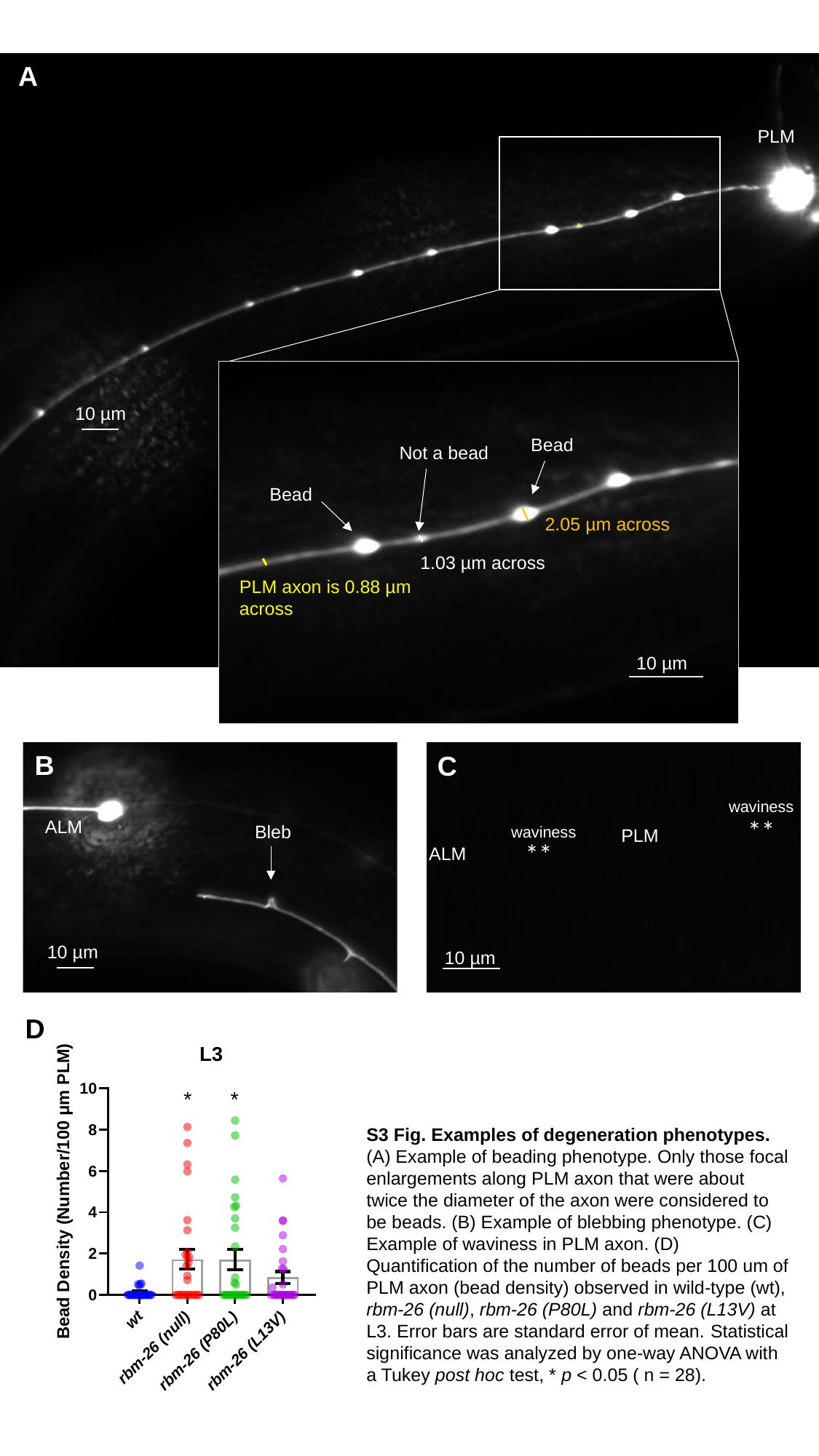

A
PLM
10 µm
Bead
Not a bead
Bead
2.05 µm across
1.03 µm across
PLM axon is 0.88 µm across
10 µm
B
B
C
waviness
**
ALM
Bleb
waviness
PLM
**
ALM
10 µm
10 µm
10 µm
D
*
*
S3 Fig. Examples of degeneration phenotypes.
(A) Example of beading phenotype. Only those focal enlargements along PLM axon that were about twice the diameter of the axon were considered to be beads. (B) Example of blebbing phenotype. (C) Example of waviness in PLM axon. (D) Quantification of the number of beads per 100 um of PLM axon (bead density) observed in wild-type (wt), rbm-26 (null), rbm-26 (P80L) and rbm-26 (L13V) at L3. Error bars are standard error of mean. Statistical significance was analyzed by one-way ANOVA with a Tukey post hoc test, * p < 0.05 ( n = 28).

### Slide 4
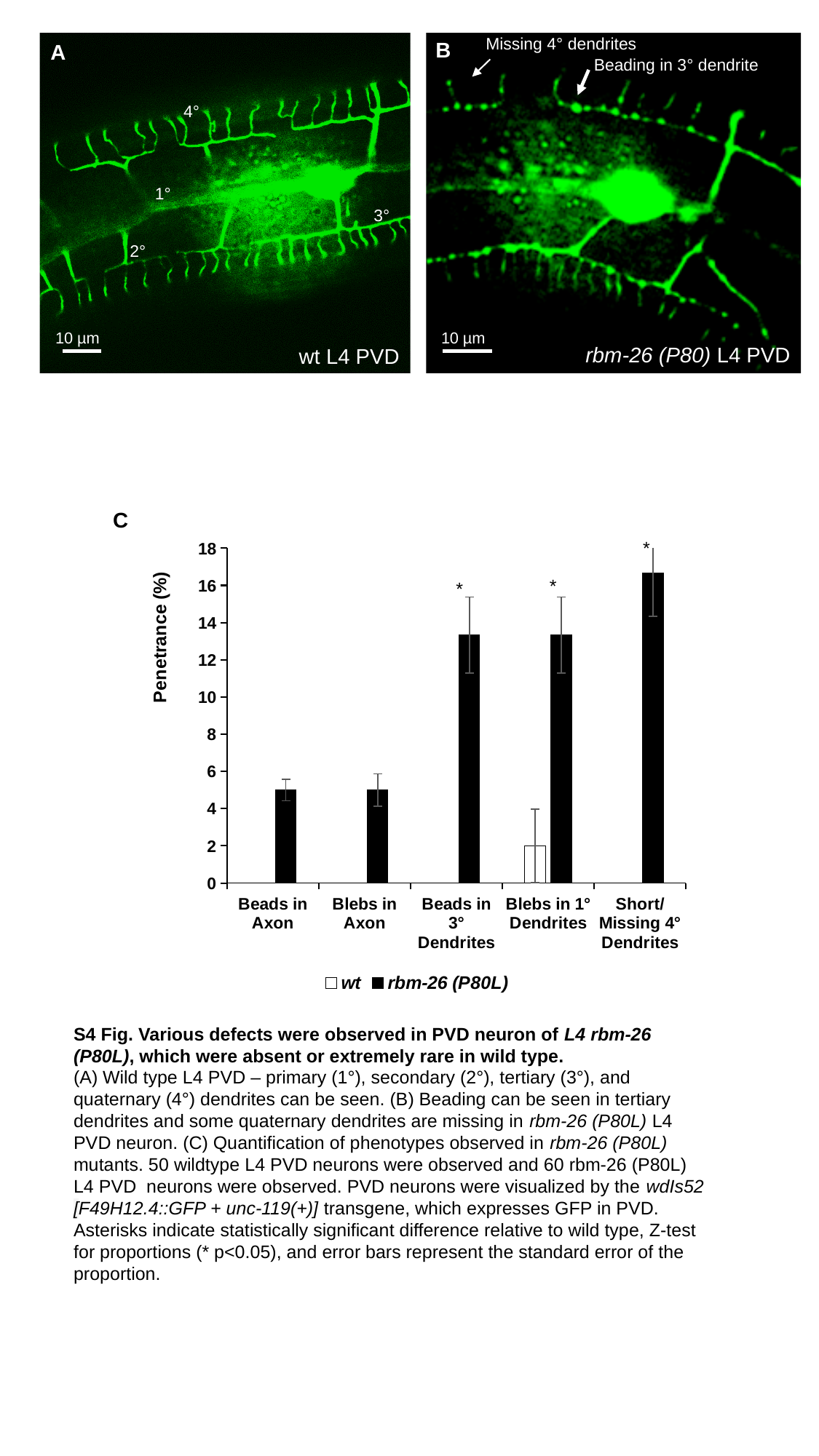

Missing 4° dendrites
B
A
Beading in 3° dendrite
4°
1°
3°
2°
10 µm
10 µm
rbm-26 (P80) L4 PVD
wt L4 PVD
C
#### Chart
| Category | wt | rbm-26 (P80L) |
|---|---|---|
| Beads in Axon | 0.0 | 5.0 |
| Blebs in Axon | 0.0 | 5.0 |
| Beads in 3° Dendrites | 0.0 | 13.333333333333334 |
| Blebs in 1° Dendrites | 2.0 | 13.333333333333334 |
| Short/Missing 4° Dendrites | 0.0 | 16.666666666666664 |*
*
*
S4 Fig. Various defects were observed in PVD neuron of L4 rbm-26 (P80L), which were absent or extremely rare in wild type.
(A) Wild type L4 PVD – primary (1°), secondary (2°), tertiary (3°), and quaternary (4°) dendrites can be seen. (B) Beading can be seen in tertiary dendrites and some quaternary dendrites are missing in rbm-26 (P80L) L4 PVD neuron. (C) Quantification of phenotypes observed in rbm-26 (P80L) mutants. 50 wildtype L4 PVD neurons were observed and 60 rbm-26 (P80L) L4 PVD neurons were observed. PVD neurons were visualized by the wdIs52 [F49H12.4::GFP + unc-119(+)] transgene, which expresses GFP in PVD. Asterisks indicate statistically significant difference relative to wild type, Z-test for proportions (* p<0.05), and error bars represent the standard error of the proportion.

### Slide 5
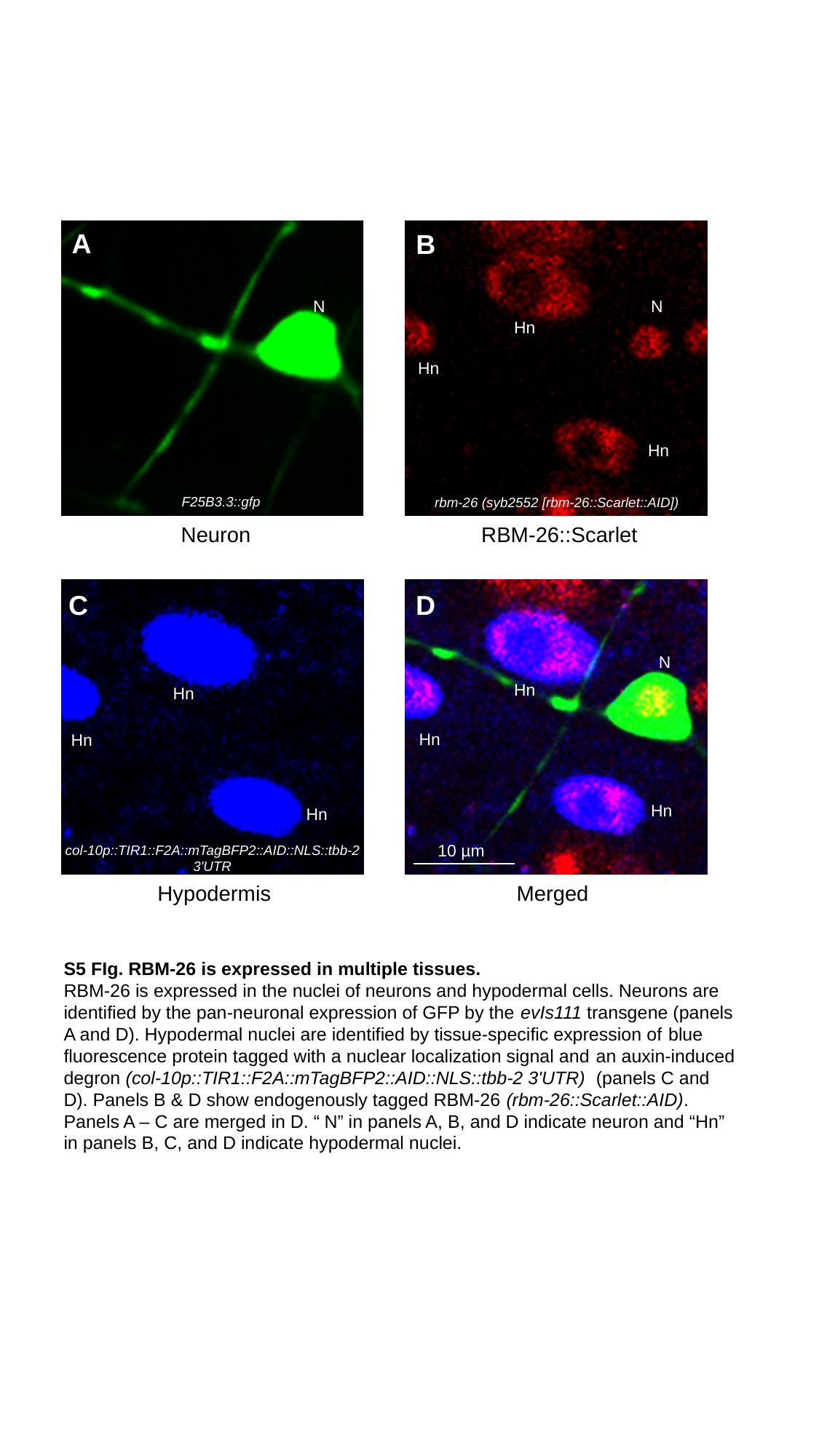

A
B
N
N
Hn
Hn
Hn
F25B3.3::gfp
rbm-26 (syb2552 [rbm-26::Scarlet::AID])
Neuron
RBM-26::Scarlet
C
D
N
Hn
Hn
Hn
Hn
Hn
Hn
10 µm
col-10p::TIR1::F2A::mTagBFP2::AID::NLS::tbb-2 3'UTR
Hypodermis
Merged
S5 FIg. RBM-26 is expressed in multiple tissues.
RBM-26 is expressed in the nuclei of neurons and hypodermal cells. Neurons are identified by the pan-neuronal expression of GFP by the evIs111 transgene (panels A and D). Hypodermal nuclei are identified by tissue-specific expression of blue fluorescence protein tagged with a nuclear localization signal and an auxin-induced degron (col-10p::TIR1::F2A::mTagBFP2::AID::NLS::tbb-2 3'UTR) (panels C and D). Panels B & D show endogenously tagged RBM-26 (rbm-26::Scarlet::AID). Panels A – C are merged in D. “ N” in panels A, B, and D indicate neuron and “Hn” in panels B, C, and D indicate hypodermal nuclei.

### Slide 6
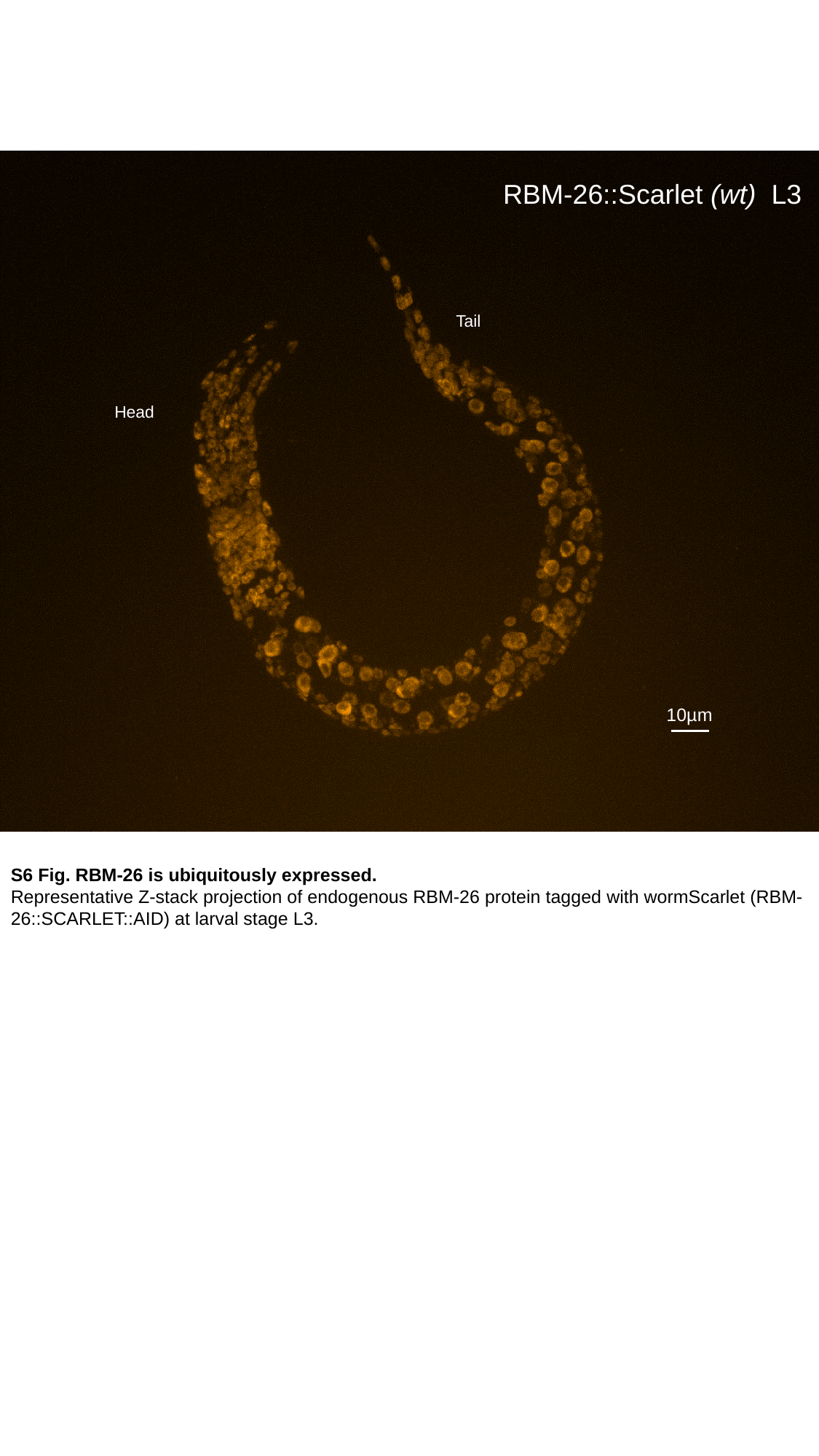

RBM-26::Scarlet (wt) L3
Tail
Head
10µm
S6 Fig. RBM-26 is ubiquitously expressed.
Representative Z-stack projection of endogenous RBM-26 protein tagged with wormScarlet (RBM-26::SCARLET::AID) at larval stage L3.

### Slide 7
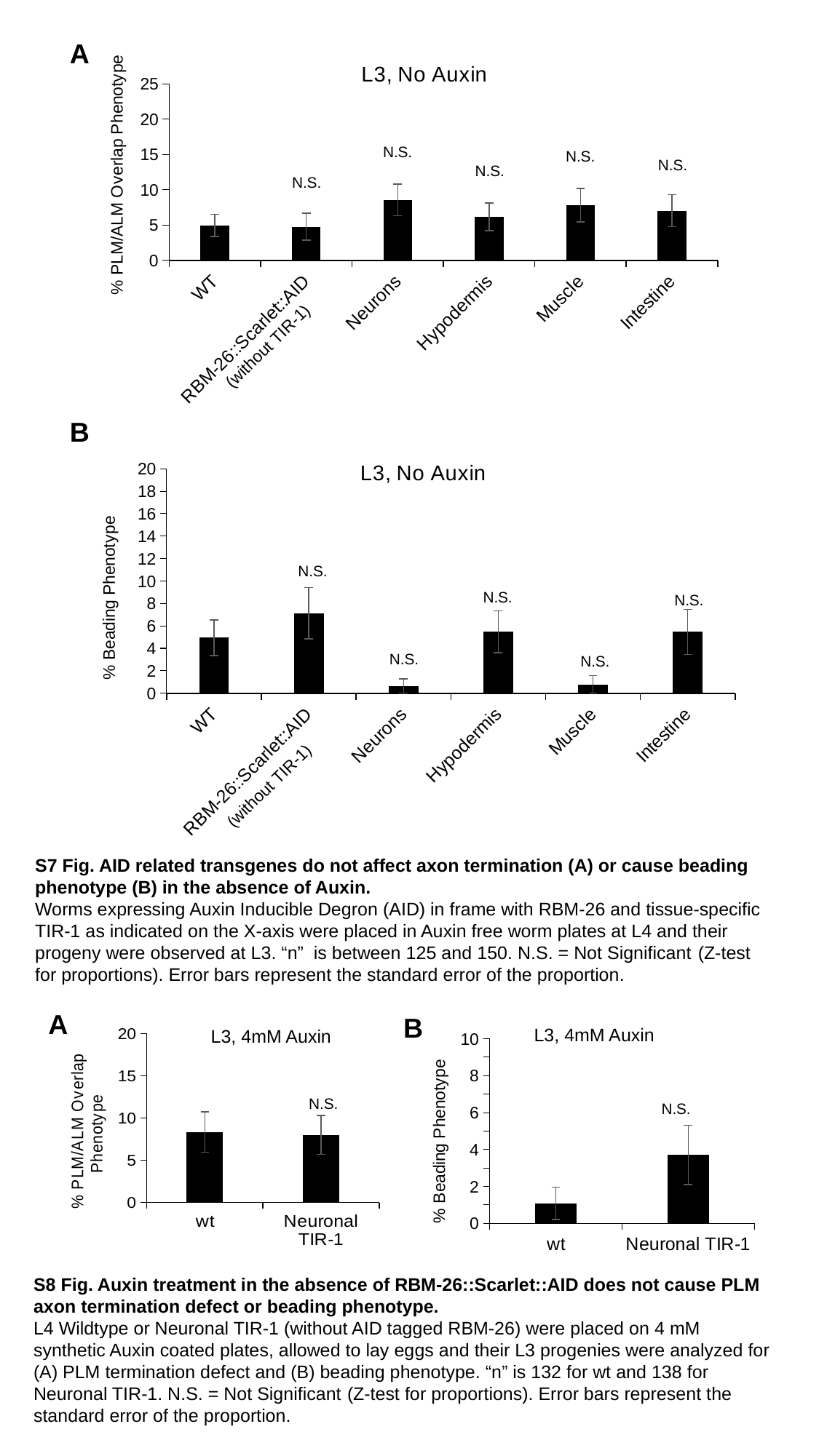

#### Chart: L3, No Auxin
| Category | PLM Overextensions |
|---|---|
| WT | 4.945054945054945 |
| RBM-26::Scarlet::AID | 4.761904761904762 |
| Neurons | 8.552631578947368 |
| Hypodermis | 6.164383561643835 |
| Muscle | 7.8125 |
| Intestine | 7.03125 |A
N.S.
N.S.
N.S.
N.S.
N.S.
(without TIR-1)
#### Chart: L3, No Auxin
| Category | PLM Beading |
|---|---|
| WT | 4.945054945054945 |
| RBM-26::Scarlet::AID | 7.142857142857142 |
| Neurons | 0.641025641025641 |
| Hypodermis | 5.47945205479452 |
| Muscle | 0.78125 |
| Intestine | 5.46875 |B
N.S.
N.S.
N.S.
N.S.
N.S.
(without TIR-1)
S7 Fig. AID related transgenes do not affect axon termination (A) or cause beading phenotype (B) in the absence of Auxin.
Worms expressing Auxin Inducible Degron (AID) in frame with RBM-26 and tissue-specific TIR-1 as indicated on the X-axis were placed in Auxin free worm plates at L4 and their progeny were observed at L3. “n” is between 125 and 150. N.S. = Not Significant (Z-test for proportions). Error bars represent the standard error of the proportion.
A
B
L3, 4mM Auxin
L3, 4mM Auxin
#### Chart
| Category | |
|---|---|
| wt | 1.0869565217391304 |
| Neuronal TIR-1 | 3.7037037037037033 |
#### Chart
| Category | |
|---|---|
| wt | 8.333333333333332 |
| Neuronal TIR-1 | 7.971014492753622 |N.S.
N.S.
S8 Fig. Auxin treatment in the absence of RBM-26::Scarlet::AID does not cause PLM axon termination defect or beading phenotype.
L4 Wildtype or Neuronal TIR-1 (without AID tagged RBM-26) were placed on 4 mM synthetic Auxin coated plates, allowed to lay eggs and their L3 progenies were analyzed for (A) PLM termination defect and (B) beading phenotype. “n” is 132 for wt and 138 for Neuronal TIR-1. N.S. = Not Significant (Z-test for proportions). Error bars represent the standard error of the proportion.

### Slide 8
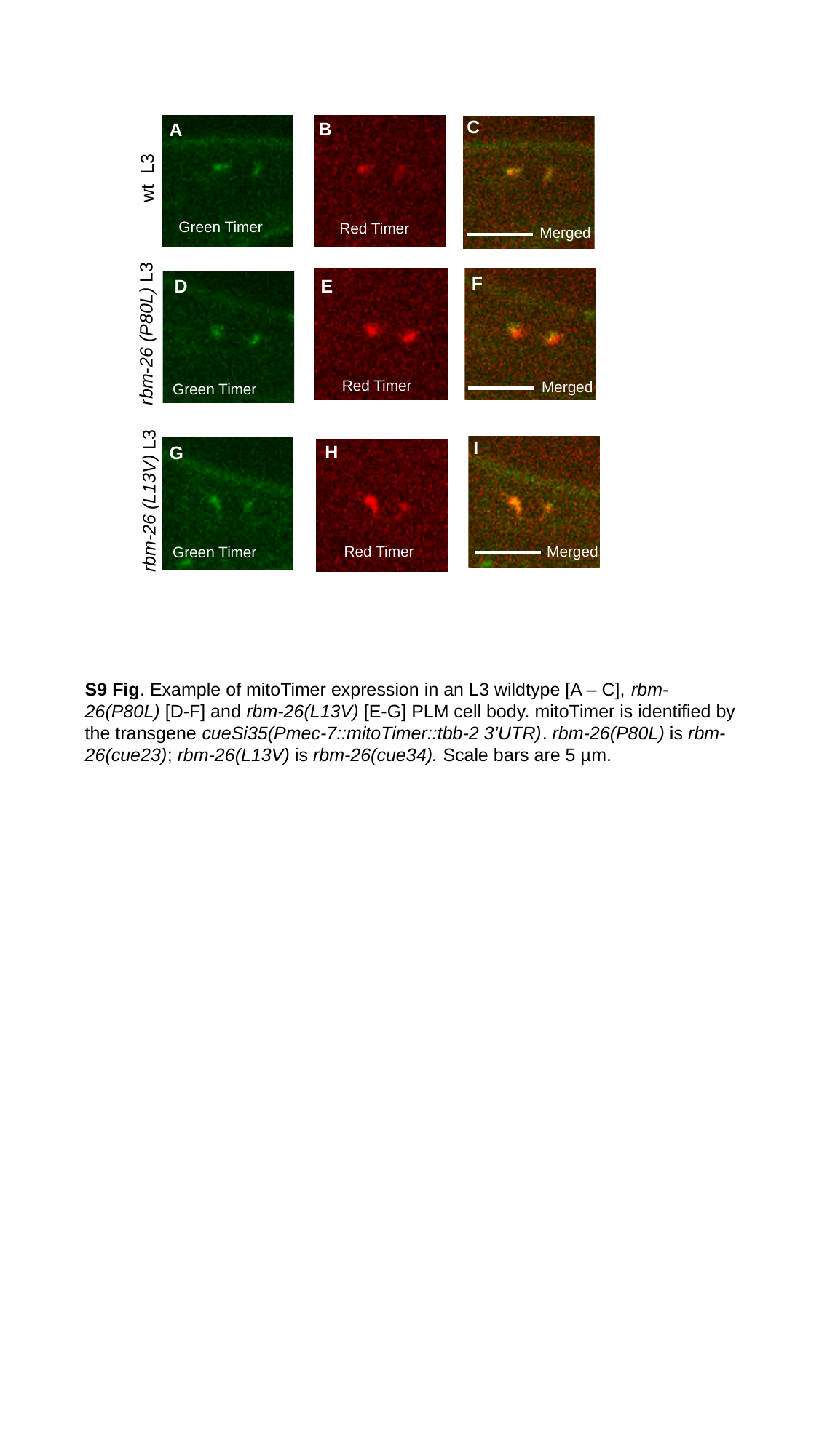

C
B
A
wt L3
Green Timer
Red Timer
Merged
F
E
D
rbm-26 (P80L) L3
Red Timer
Merged
Green Timer
I
H
G
rbm-26 (L13V) L3
Red Timer
Merged
Green Timer
S9 Fig. Example of mitoTimer expression in an L3 wildtype [A – C], rbm-26(P80L) [D-F] and rbm-26(L13V) [E-G] PLM cell body. mitoTimer is identified by the transgene cueSi35(Pmec-7::mitoTimer::tbb-2 3’UTR). rbm-26(P80L) is rbm-26(cue23); rbm-26(L13V) is rbm-26(cue34). Scale bars are 5 µm.

### Slide 9
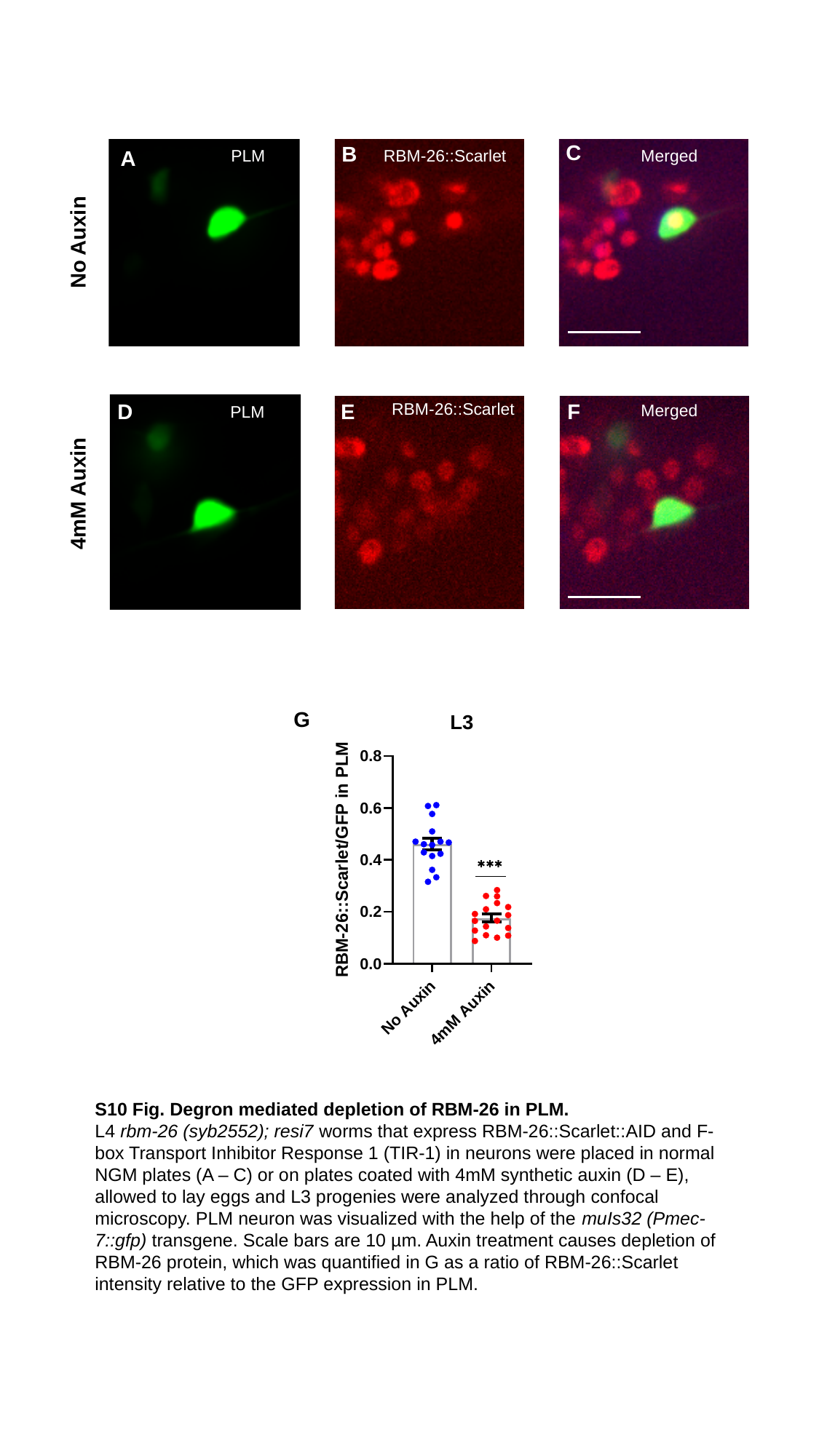

C
B
No Auxin
A
PLM
RBM-26::Scarlet
Merged
4mM Auxin
E
F
D
RBM-26::Scarlet
Merged
PLM
G
S10 Fig. Degron mediated depletion of RBM-26 in PLM.
L4 rbm-26 (syb2552); resi7 worms that express RBM-26::Scarlet::AID and F-box Transport Inhibitor Response 1 (TIR-1) in neurons were placed in normal NGM plates (A – C) or on plates coated with 4mM synthetic auxin (D – E), allowed to lay eggs and L3 progenies were analyzed through confocal microscopy. PLM neuron was visualized with the help of the muIs32 (Pmec-7::gfp) transgene. Scale bars are 10 µm. Auxin treatment causes depletion of RBM-26 protein, which was quantified in G as a ratio of RBM-26::Scarlet intensity relative to the GFP expression in PLM.

### Slide 10
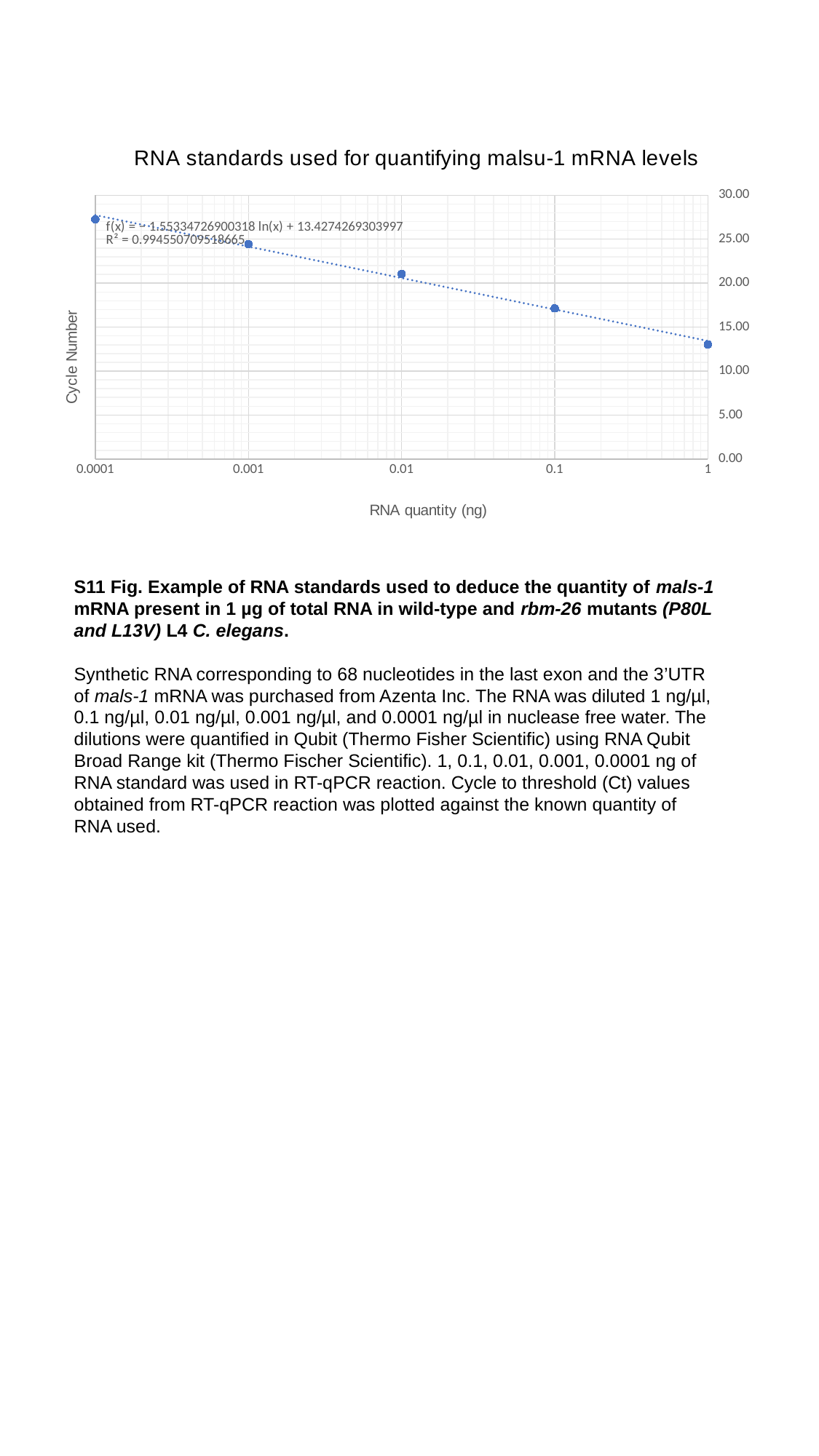

#### Chart: RNA standards used for quantifying malsu-1 mRNA levels
| Category | |
|---|---|S11 Fig. Example of RNA standards used to deduce the quantity of mals-1 mRNA present in 1 µg of total RNA in wild-type and rbm-26 mutants (P80L and L13V) L4 C. elegans.
Synthetic RNA corresponding to 68 nucleotides in the last exon and the 3’UTR of mals-1 mRNA was purchased from Azenta Inc. The RNA was diluted 1 ng/µl, 0.1 ng/µl, 0.01 ng/µl, 0.001 ng/µl, and 0.0001 ng/µl in nuclease free water. The dilutions were quantified in Qubit (Thermo Fisher Scientific) using RNA Qubit Broad Range kit (Thermo Fischer Scientific). 1, 0.1, 0.01, 0.001, 0.0001 ng of RNA standard was used in RT-qPCR reaction. Cycle to threshold (Ct) values obtained from RT-qPCR reaction was plotted against the known quantity of RNA used.

### Slide 11
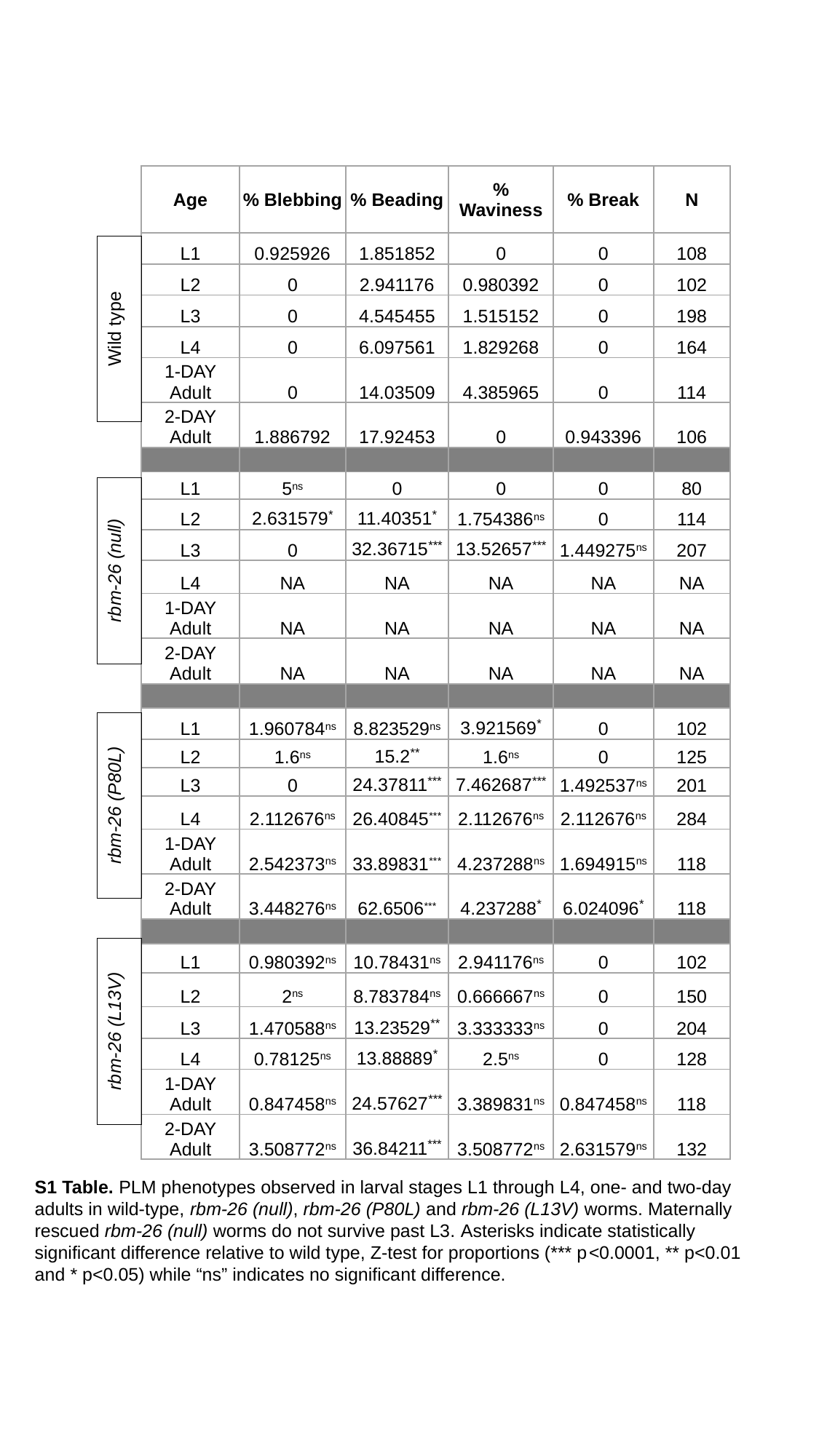

| Age | % Blebbing | % Beading | % Waviness | % Break | N |
| --- | --- | --- | --- | --- | --- |
| L1 | 0.925926 | 1.851852 | 0 | 0 | 108 |
| L2 | 0 | 2.941176 | 0.980392 | 0 | 102 |
| L3 | 0 | 4.545455 | 1.515152 | 0 | 198 |
| L4 | 0 | 6.097561 | 1.829268 | 0 | 164 |
| 1-DAY Adult | 0 | 14.03509 | 4.385965 | 0 | 114 |
| 2-DAY Adult | 1.886792 | 17.92453 | 0 | 0.943396 | 106 |
| L1 | 5^ns^ | 0 | 0 | 0 | 80 |
| L2 | 2.631579^\*^ | 11.40351^\*^ | 1.754386^ns^ | 0 | 114 |
| L3 | 0 | 32.36715^\*\*\*^ | 13.52657^\*\*\*^ | 1.449275^ns^ | 207 |
| L4 | NA | NA | NA | NA | NA |
| 1-DAY Adult | NA | NA | NA | NA | NA |
| 2-DAY Adult | NA | NA | NA | NA | NA |
| L1 | 1.960784^ns^ | 8.823529^ns^ | 3.921569^\*^ | 0 | 102 |
| L2 | 1.6^ns^ | 15.2^\*\*^ | 1.6^ns^ | 0 | 125 |
| L3 | 0 | 24.37811^\*\*\*^ | 7.462687^\*\*\*^ | 1.492537^ns^ | 201 |
| L4 | 2.112676^ns^ | 26.40845^\*\*\*^ | 2.112676^ns^ | 2.112676^ns^ | 284 |
| 1-DAY Adult | 2.542373^ns^ | 33.89831^\*\*\*^ | 4.237288^ns^ | 1.694915^ns^ | 118 |
| 2-DAY Adult | 3.448276^ns^ | 62.6506^\*\*\*^ | 4.237288^\*^ | 6.024096^\*^ | 118 |
| L1 | 0.980392^ns^ | 10.78431^ns^ | 2.941176^ns^ | 0 | 102 |
| L2 | 2^ns^ | 8.783784^ns^ | 0.666667^ns^ | 0 | 150 |
| L3 | 1.470588^ns^ | 13.23529^\*\*^ | 3.333333^ns^ | 0 | 204 |
| L4 | 0.78125^ns^ | 13.88889^\*^ | 2.5^ns^ | 0 | 128 |
| 1-DAY Adult | 0.847458^ns^ | 24.57627^\*\*\*^ | 3.389831^ns^ | 0.847458^ns^ | 118 |
| 2-DAY Adult | 3.508772^ns^ | 36.84211^\*\*\*^ | 3.508772^ns^ | 2.631579^ns^ | 132 |
Wild type
rbm-26 (null)
rbm-26 (P80L)
rbm-26 (L13V)
S1 Table. PLM phenotypes observed in larval stages L1 through L4, one- and two-day adults in wild-type, rbm-26 (null), rbm-26 (P80L) and rbm-26 (L13V) worms. Maternally rescued rbm-26 (null) worms do not survive past L3. Asterisks indicate statistically significant difference relative to wild type, Z-test for proportions (*** p<0.0001, ** p<0.01 and * p<0.05) while “ns” indicates no significant difference.

### Slide 12
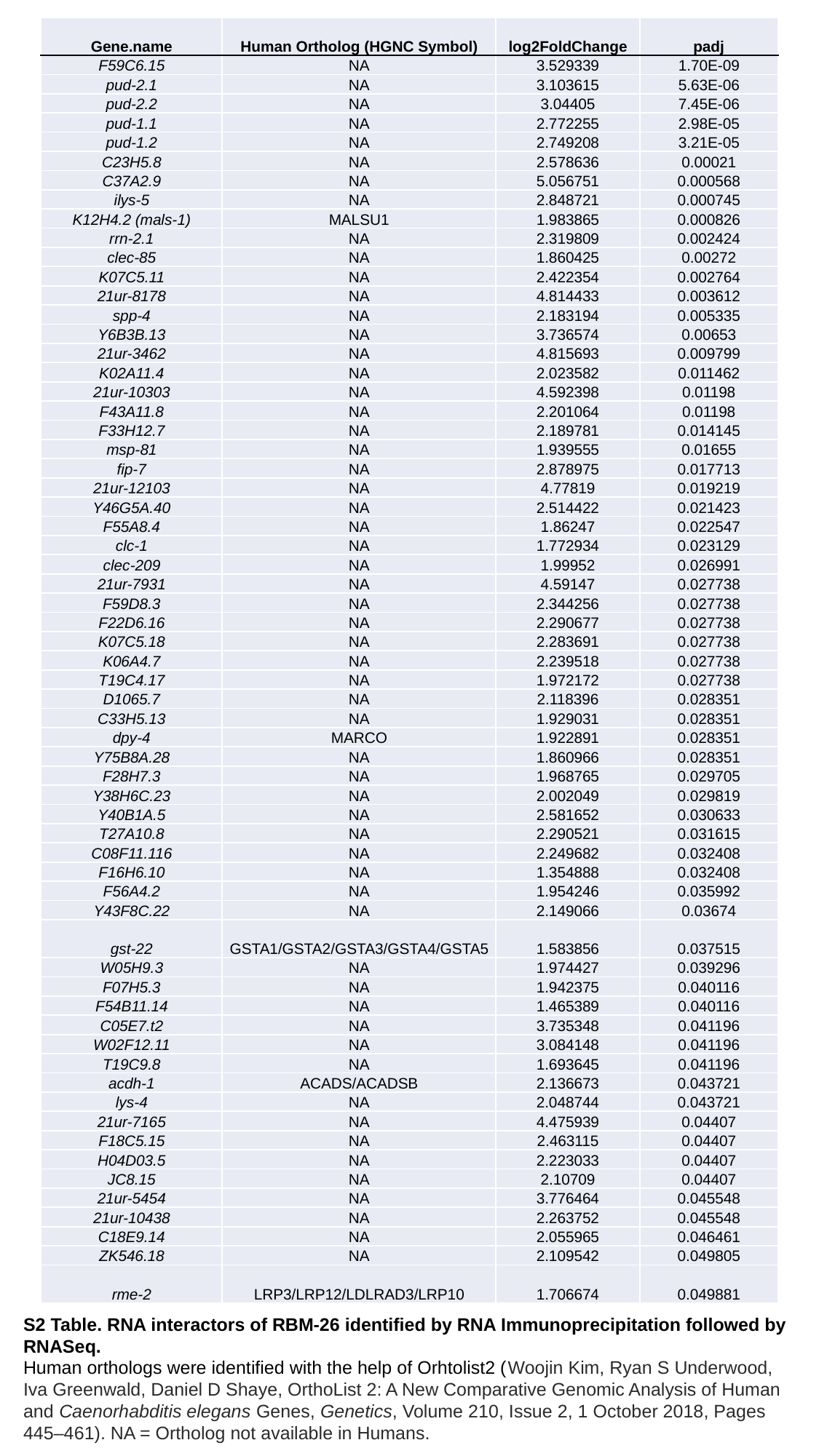

| Gene.name | Human Ortholog (HGNC Symbol) | log2FoldChange | padj |
| --- | --- | --- | --- |
| F59C6.15 | NA | 3.529339 | 1.70E-09 |
| pud-2.1 | NA | 3.103615 | 5.63E-06 |
| pud-2.2 | NA | 3.04405 | 7.45E-06 |
| pud-1.1 | NA | 2.772255 | 2.98E-05 |
| pud-1.2 | NA | 2.749208 | 3.21E-05 |
| C23H5.8 | NA | 2.578636 | 0.00021 |
| C37A2.9 | NA | 5.056751 | 0.000568 |
| ilys-5 | NA | 2.848721 | 0.000745 |
| K12H4.2 (mals-1) | MALSU1 | 1.983865 | 0.000826 |
| rrn-2.1 | NA | 2.319809 | 0.002424 |
| clec-85 | NA | 1.860425 | 0.00272 |
| K07C5.11 | NA | 2.422354 | 0.002764 |
| 21ur-8178 | NA | 4.814433 | 0.003612 |
| spp-4 | NA | 2.183194 | 0.005335 |
| Y6B3B.13 | NA | 3.736574 | 0.00653 |
| 21ur-3462 | NA | 4.815693 | 0.009799 |
| K02A11.4 | NA | 2.023582 | 0.011462 |
| 21ur-10303 | NA | 4.592398 | 0.01198 |
| F43A11.8 | NA | 2.201064 | 0.01198 |
| F33H12.7 | NA | 2.189781 | 0.014145 |
| msp-81 | NA | 1.939555 | 0.01655 |
| fip-7 | NA | 2.878975 | 0.017713 |
| 21ur-12103 | NA | 4.77819 | 0.019219 |
| Y46G5A.40 | NA | 2.514422 | 0.021423 |
| F55A8.4 | NA | 1.86247 | 0.022547 |
| clc-1 | NA | 1.772934 | 0.023129 |
| clec-209 | NA | 1.99952 | 0.026991 |
| 21ur-7931 | NA | 4.59147 | 0.027738 |
| F59D8.3 | NA | 2.344256 | 0.027738 |
| F22D6.16 | NA | 2.290677 | 0.027738 |
| K07C5.18 | NA | 2.283691 | 0.027738 |
| K06A4.7 | NA | 2.239518 | 0.027738 |
| T19C4.17 | NA | 1.972172 | 0.027738 |
| D1065.7 | NA | 2.118396 | 0.028351 |
| C33H5.13 | NA | 1.929031 | 0.028351 |
| dpy-4 | MARCO | 1.922891 | 0.028351 |
| Y75B8A.28 | NA | 1.860966 | 0.028351 |
| F28H7.3 | NA | 1.968765 | 0.029705 |
| Y38H6C.23 | NA | 2.002049 | 0.029819 |
| Y40B1A.5 | NA | 2.581652 | 0.030633 |
| T27A10.8 | NA | 2.290521 | 0.031615 |
| C08F11.116 | NA | 2.249682 | 0.032408 |
| F16H6.10 | NA | 1.354888 | 0.032408 |
| F56A4.2 | NA | 1.954246 | 0.035992 |
| Y43F8C.22 | NA | 2.149066 | 0.03674 |
| gst-22 | GSTA1/GSTA2/GSTA3/GSTA4/GSTA5 | 1.583856 | 0.037515 |
| W05H9.3 | NA | 1.974427 | 0.039296 |
| F07H5.3 | NA | 1.942375 | 0.040116 |
| F54B11.14 | NA | 1.465389 | 0.040116 |
| C05E7.t2 | NA | 3.735348 | 0.041196 |
| W02F12.11 | NA | 3.084148 | 0.041196 |
| T19C9.8 | NA | 1.693645 | 0.041196 |
| acdh-1 | ACADS/ACADSB | 2.136673 | 0.043721 |
| lys-4 | NA | 2.048744 | 0.043721 |
| 21ur-7165 | NA | 4.475939 | 0.04407 |
| F18C5.15 | NA | 2.463115 | 0.04407 |
| H04D03.5 | NA | 2.223033 | 0.04407 |
| JC8.15 | NA | 2.10709 | 0.04407 |
| 21ur-5454 | NA | 3.776464 | 0.045548 |
| 21ur-10438 | NA | 2.263752 | 0.045548 |
| C18E9.14 | NA | 2.055965 | 0.046461 |
| ZK546.18 | NA | 2.109542 | 0.049805 |
| rme-2 | LRP3/LRP12/LDLRAD3/LRP10 | 1.706674 | 0.049881 |
S2 Table. RNA interactors of RBM-26 identified by RNA Immunoprecipitation followed by RNASeq.
Human orthologs were identified with the help of Orhtolist2 (Woojin Kim, Ryan S Underwood, Iva Greenwald, Daniel D Shaye, OrthoList 2: A New Comparative Genomic Analysis of Human and Caenorhabditis elegans Genes, Genetics, Volume 210, Issue 2, 1 October 2018, Pages 445–461). NA = Ortholog not available in Humans.

### Slide 13
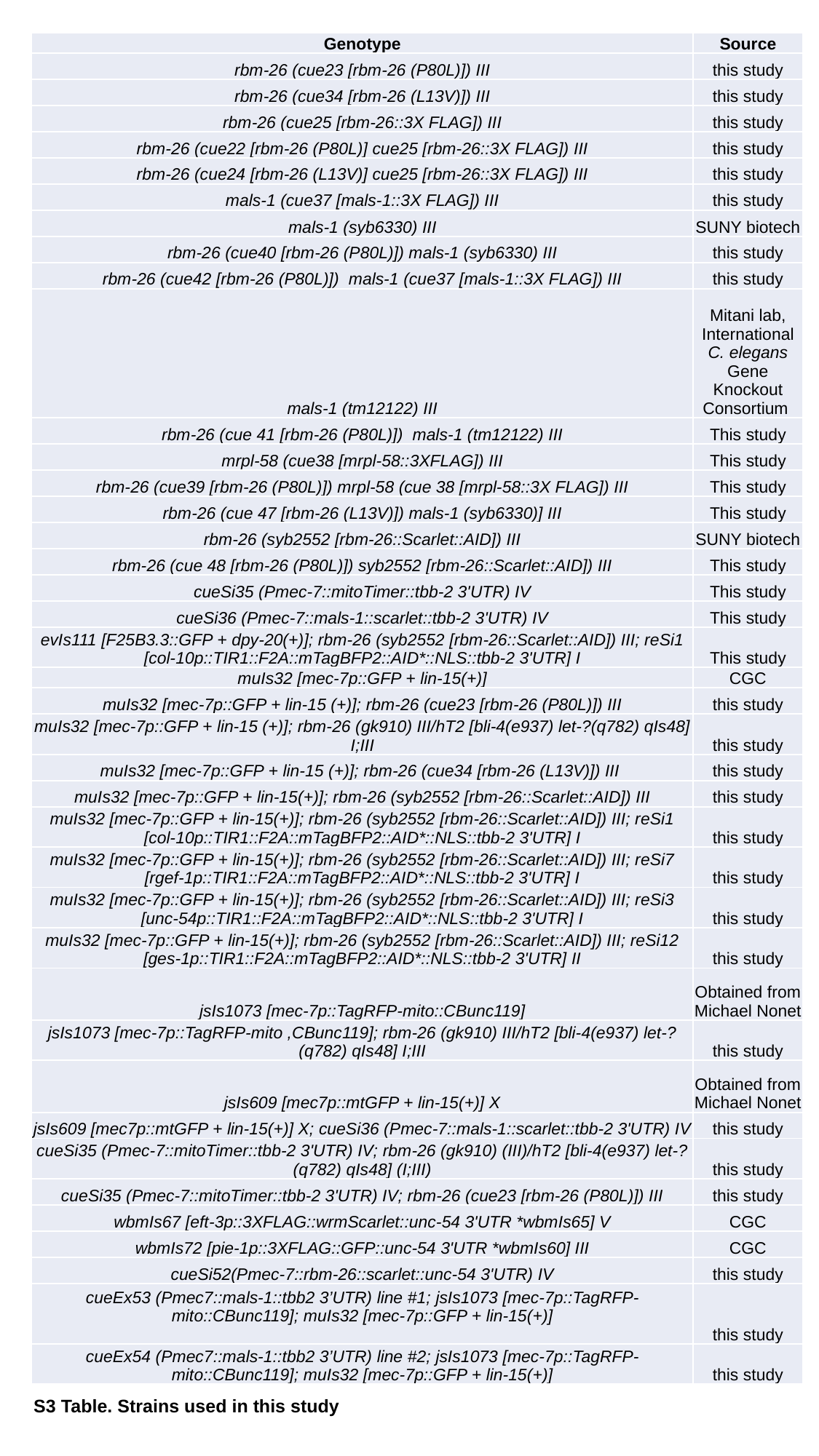

| Genotype | Source |
| --- | --- |
| rbm-26 (cue23 [rbm-26 (P80L)]) III | this study |
| rbm-26 (cue34 [rbm-26 (L13V)]) III | this study |
| rbm-26 (cue25 [rbm-26::3X FLAG]) III | this study |
| rbm-26 (cue22 [rbm-26 (P80L)] cue25 [rbm-26::3X FLAG]) III | this study |
| rbm-26 (cue24 [rbm-26 (L13V)] cue25 [rbm-26::3X FLAG]) III | this study |
| mals-1 (cue37 [mals-1::3X FLAG]) III | this study |
| mals-1 (syb6330) III | SUNY biotech |
| rbm-26 (cue40 [rbm-26 (P80L)]) mals-1 (syb6330) III | this study |
| rbm-26 (cue42 [rbm-26 (P80L)]) mals-1 (cue37 [mals-1::3X FLAG]) III | this study |
| mals-1 (tm12122) III | Mitani lab, International C. elegans Gene Knockout Consortium |
| rbm-26 (cue 41 [rbm-26 (P80L)]) mals-1 (tm12122) III | This study |
| mrpl-58 (cue38 [mrpl-58::3XFLAG]) III | This study |
| rbm-26 (cue39 [rbm-26 (P80L)]) mrpl-58 (cue 38 [mrpl-58::3X FLAG]) III | This study |
| rbm-26 (cue 47 [rbm-26 (L13V)]) mals-1 (syb6330)] III | This study |
| rbm-26 (syb2552 [rbm-26::Scarlet::AID]) III | SUNY biotech |
| rbm-26 (cue 48 [rbm-26 (P80L)]) syb2552 [rbm-26::Scarlet::AID]) III | This study |
| cueSi35 (Pmec-7::mitoTimer::tbb-2 3'UTR) IV | This study |
| cueSi36 (Pmec-7::mals-1::scarlet::tbb-2 3'UTR) IV | This study |
| evIs111 [F25B3.3::GFP + dpy-20(+)]; rbm-26 (syb2552 [rbm-26::Scarlet::AID]) III; reSi1 [col-10p::TIR1::F2A::mTagBFP2::AID\*::NLS::tbb-2 3'UTR] I | This study |
| muIs32 [mec-7p::GFP + lin-15(+)] | CGC |
| muIs32 [mec-7p::GFP + lin-15 (+)]; rbm-26 (cue23 [rbm-26 (P80L)]) III | this study |
| muIs32 [mec-7p::GFP + lin-15 (+)]; rbm-26 (gk910) III/hT2 [bli-4(e937) let-?(q782) qIs48] I;III | this study |
| muIs32 [mec-7p::GFP + lin-15 (+)]; rbm-26 (cue34 [rbm-26 (L13V)]) III | this study |
| muIs32 [mec-7p::GFP + lin-15(+)]; rbm-26 (syb2552 [rbm-26::Scarlet::AID]) III | this study |
| muIs32 [mec-7p::GFP + lin-15(+)]; rbm-26 (syb2552 [rbm-26::Scarlet::AID]) III; reSi1 [col-10p::TIR1::F2A::mTagBFP2::AID\*::NLS::tbb-2 3'UTR] I | this study |
| muIs32 [mec-7p::GFP + lin-15(+)]; rbm-26 (syb2552 [rbm-26::Scarlet::AID]) III; reSi7 [rgef-1p::TIR1::F2A::mTagBFP2::AID\*::NLS::tbb-2 3'UTR] I | this study |
| muIs32 [mec-7p::GFP + lin-15(+)]; rbm-26 (syb2552 [rbm-26::Scarlet::AID]) III; reSi3 [unc-54p::TIR1::F2A::mTagBFP2::AID\*::NLS::tbb-2 3'UTR] I | this study |
| muIs32 [mec-7p::GFP + lin-15(+)]; rbm-26 (syb2552 [rbm-26::Scarlet::AID]) III; reSi12 [ges-1p::TIR1::F2A::mTagBFP2::AID\*::NLS::tbb-2 3'UTR] II | this study |
| jsIs1073 [mec-7p::TagRFP-mito::CBunc119] | Obtained from Michael Nonet |
| jsIs1073 [mec-7p::TagRFP-mito ,CBunc119]; rbm-26 (gk910) III/hT2 [bli-4(e937) let-?(q782) qIs48] I;III | this study |
| jsIs609 [mec7p::mtGFP + lin-15(+)] X | Obtained from Michael Nonet |
| jsIs609 [mec7p::mtGFP + lin-15(+)] X; cueSi36 (Pmec-7::mals-1::scarlet::tbb-2 3'UTR) IV | this study |
| cueSi35 (Pmec-7::mitoTimer::tbb-2 3'UTR) IV; rbm-26 (gk910) (III)/hT2 [bli-4(e937) let-?(q782) qIs48] (I;III) | this study |
| cueSi35 (Pmec-7::mitoTimer::tbb-2 3'UTR) IV; rbm-26 (cue23 [rbm-26 (P80L)]) III | this study |
| wbmIs67 [eft-3p::3XFLAG::wrmScarlet::unc-54 3'UTR \*wbmIs65] V | CGC |
| wbmIs72 [pie-1p::3XFLAG::GFP::unc-54 3'UTR \*wbmIs60] III | CGC |
| cueSi52(Pmec-7::rbm-26::scarlet::unc-54 3'UTR) IV | this study |
| cueEx53 (Pmec7::mals-1::tbb2 3’UTR) line #1; jsIs1073 [mec-7p::TagRFP-mito::CBunc119]; muIs32 [mec-7p::GFP + lin-15(+)] | this study |
| cueEx54 (Pmec7::mals-1::tbb2 3’UTR) line #2; jsIs1073 [mec-7p::TagRFP-mito::CBunc119]; muIs32 [mec-7p::GFP + lin-15(+)] | this study |
S3 Table. Strains used in this study

### Slide 14
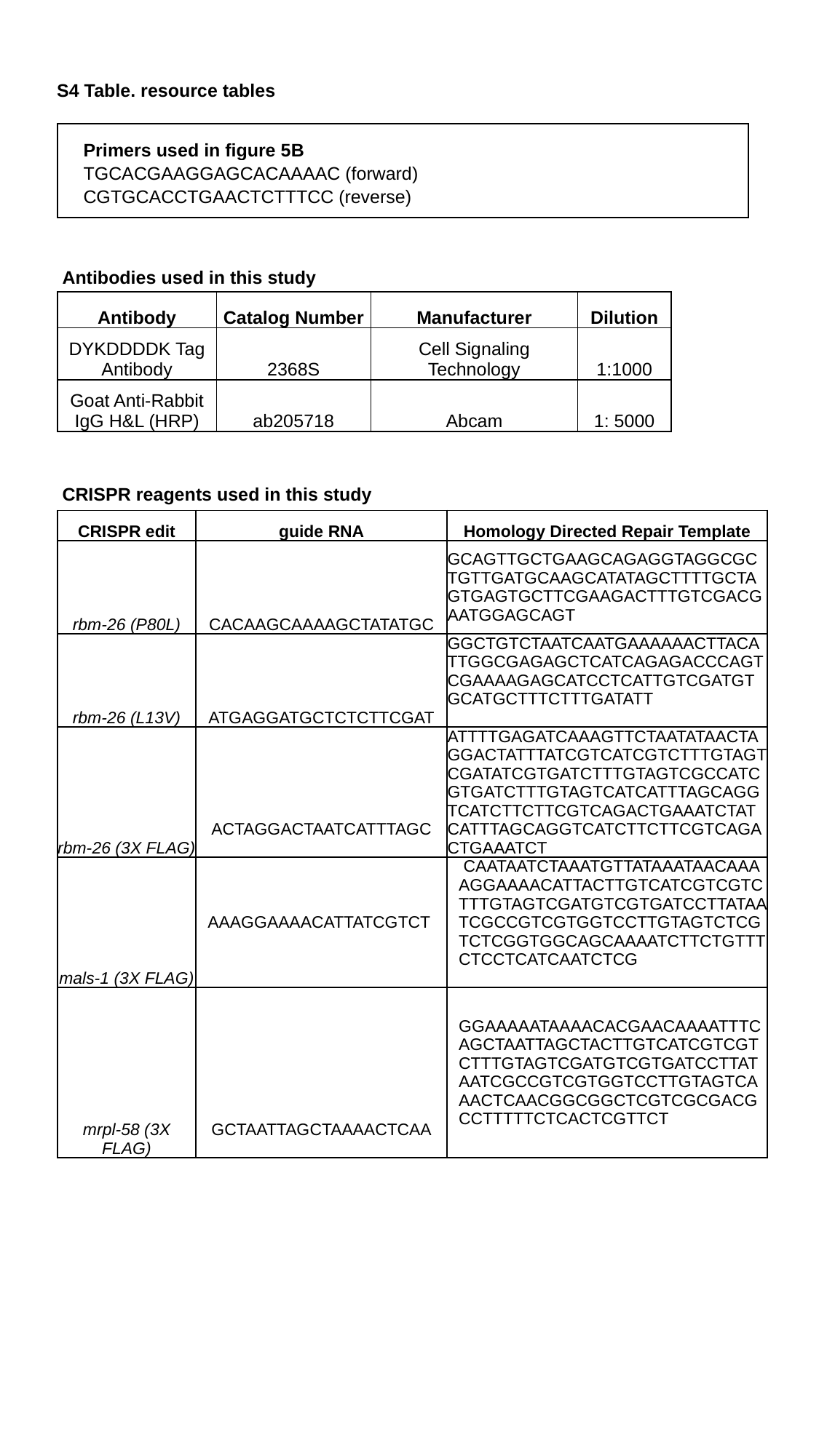

S4 Table. resource tables
| Primers used in figure 5B |
| --- |
| TGCACGAAGGAGCACAAAAC (forward) |
| CGTGCACCTGAACTCTTTCC (reverse) |
Antibodies used in this study
| Antibody | Catalog Number | Manufacturer | Dilution |
| --- | --- | --- | --- |
| DYKDDDDK Tag Antibody | 2368S | Cell Signaling Technology | 1:1000 |
| Goat Anti-Rabbit IgG H&L (HRP) | ab205718 | Abcam | 1: 5000 |
CRISPR reagents used in this study
| CRISPR edit | guide RNA | Homology Directed Repair Template |
| --- | --- | --- |
| rbm-26 (P80L) | CACAAGCAAAAGCTATATGC | GCAGTTGCTGAAGCAGAGGTAGGCGCTGTTGATGCAAGCATATAGCTTTTGCTAGTGAGTGCTTCGAAGACTTTGTCGACGAATGGAGCAGT |
| rbm-26 (L13V) | ATGAGGATGCTCTCTTCGAT | GGCTGTCTAATCAATGAAAAAACTTACATTGGCGAGAGCTCATCAGAGACCCAGTCGAAAAGAGCATCCTCATTGTCGATGTGCATGCTTTCTTTGATATT |
| rbm-26 (3X FLAG) | ACTAGGACTAATCATTTAGC | ATTTTGAGATCAAAGTTCTAATATAACTAGGACTATTTATCGTCATCGTCTTTGTAGTCGATATCGTGATCTTTGTAGTCGCCATCGTGATCTTTGTAGTCATCATTTAGCAGGTCATCTTCTTCGTCAGACTGAAATCTATCATTTAGCAGGTCATCTTCTTCGTCAGACTGAAATCT |
| mals-1 (3X FLAG) | AAAGGAAAACATTATCGTCT | CAATAATCTAAATGTTATAAATAACAAAAGGAAAACATTACTTGTCATCGTCGTCTTTGTAGTCGATGTCGTGATCCTTATAATCGCCGTCGTGGTCCTTGTAGTCTCGTCTCGGTGGCAGCAAAATCTTCTGTTTCTCCTCATCAATCTCG |
| mrpl-58 (3X FLAG) | GCTAATTAGCTAAAACTCAA | GGAAAAATAAAACACGAACAAAATTTCAGCTAATTAGCTACTTGTCATCGTCGTCTTTGTAGTCGATGTCGTGATCCTTATAATCGCCGTCGTGGTCCTTGTAGTCAAACTCAACGGCGGCTCGTCGCGACGCCTTTTTCTCACTCGTTCT |
